# Integrating GlycoSHIELD Modeling and DNA-PAINT SMLM to Map the Glycosylation-Dependent Distribution of the Na,K-ATPase

**DOI:** 10.64898/2026.03.27.714919

**Authors:** Bruno Stojcic, Piotr Draczkowski, Joan Patrick, Mezida Saeed, Hjalmar Brismar

## Abstract

The cell surface localization of the Na,K-ATPase (sodium pump) is required for maintaining transmembrane electrochemical gradients. While glycosylation of the β1 subunit facilitates trafficking from the endoplasmic reticulum to the plasma membrane, its role in nanoscale surface organization is not characterized. This study employed GlycoSHIELD computational modeling and DNA-PAINT single-molecule localization microscopy (SMLM) to evaluate how N-glycans influence pump distribution. In-silico simulations indicated that N-glycans sequester the protein core, providing a steric shield that increases with structural complexity. To investigate this experimentally, glycosylation-deficient mutants (3NQ) were generated and confirmed via immunoblotting. Quantitative SMLM analysis of A498 cells demonstrated that wild-type pumps exhibit higher localization density and form larger (144 nm) and more frequent clusters than 3NQ mutants (109 nm). These results indicate that N-glycosylation promotes stable enzyme clustering, supporting a galectin-lattice mechanism of organization rather than steric repulsion.

## Introduction

The Na,K-ATPase (sodium pump) is an essential membrane-bound enzyme that maintains the electrochemical gradients of sodium and potassium ions across the plasma membrane. By actively exchanging three intracellular sodium ions for two extracellular potassium ions using energy from ATP hydrolysis, the pump sustains several central cellular functions including membrane potential, fluid balance regulation, nerve impulse transmission, and muscle contraction (Vagin et al. 2012; Kaplan 2002; Yordy and Bowen 1993; Jorgensen et al. 2003; Cereijido et al. 2012).

Na,K-ATPase functions as a heterotrimeric complex composed of alpha (α), beta (β), and gamma (γ) subunits. The α-subunit serves as the catalytic core of the enzyme and exists in four isoforms (α1–α4), which display tissue-specific expression patterns. This subunit contains ten transmembrane helices that form the pathway for ion transport and house the binding sites for ATP and ouabain, a known inhibitor of Na,K-ATPase. The small γ-subunit, member of the FXYD protein family, modulates the pump’s activity by affecting ion affinity and pump kinetics in an isoform-specific manner, allowing for fine-tuning to meet the physiological demands of different tissues (Kaplan 2002).

The β-subunit, with its three isoforms (β1–β3), is a highly glycosylated type II membrane protein that plays a pivotal role in the proper folding, assembly, and stabilization of the α-subunit (Ackermann and Geering 1990). Glycosylation, the enzymatic addition of carbohydrates to proteins, occurs as both a co-translational and a post-translational modification, introducing structural changes such as alterations in charge, molecular weight, and interaction partners. It also aids in proper protein folding and facilitates their transport through the endo-lysosomal pathway. Broadly, proteins are glycosylated at N-linked (asparagine) and O-linked (serine/threonine) residues.

Beyond protein folding, β-subunit is instrumental in determining the enzyme’s spatial localization within polarized cells. For instance, the β2 isoform promotes apical delivery by clustering in lipid rafts and caveolae, overriding the basolateral sorting signals of the α1 subunit. Additionally, glycosylation is vital for cell-cell adhesion where the β1 subunit associates strongly with adherent junctions and the cytoskeleton to maintain epithelial integrity. Unglycosylated β1 subunits, as seen in MDCK cells, exhibit weaker adhesion and impaired formation of cell contacts, underscoring the role of N-glycans in stabilizing intercellular junctions (Vagin et al. 2007).

While glycosylation of the β-subunit is known to be essential for the trafficking of Na,K-ATPase from the endoplasmic reticulum (ER) to the plasma membrane, its role in the spatial organization of the pump at the cell surface remains less understood. In mammalian systems, glycosylation-deficient mutants often exhibit increased retention in the ER due to quality control mechanisms that recognize them as misfolded. However, the β1 subunit can escape this mechanism and unglycosylated β-subunits can assemble with α-subunits and retain normal ion transport activity and ouabain binding (Beggah et al. 1997).This suggests that glycosylation primarily serves structural maturation and trafficking rather than directly influencing catalytic function.

A central paradox of Na,K-ATPase organization is that while it must be mobile enough to cycle through conformations, it is often found concentrated in specific membrane domains (Reinhard et al. 2013; Vagin et al. 2012). Two competing hypotheses explain how N-glycosylation on the β1 subunit influences this spatial arrangement. The Galectin-Lattice hypothesis posits that glycans act as “handles” for multivalent carbohydrate-binding proteins such as Galectin-1 or Galectin-3 (Shafaq-Zadah et al. 2025). These proteins can cross-link the N-glycans of adjacent pumps, promoting the formation of larger, denser clusters that are stabilized at the cell surface. Conversely, the Steric Repulsion hypothesis suggests that the large, highly hydrated N-glycan chains act as entropic molecular spacers. In this model, the “sugar canopy” generated by the β1 isoform constitutes a physical barrier to tight packing, preventing overcrowding that could otherwise stall the catalytic cycle.

To investigate the lateral distribution and nanoscale architecture of the Na,K-ATPase, we used Total Internal Reflection Fluorescence (TIRF)-based Single-Molecule Localization Microscopy (SMLM). Unlike conventional light microscopy, which is limited by the diffraction of light, SMLM provides the necessary spatial resolution to resolve and quantify the clustering, density, and distribution of individual enzyme complexes on the plasma membrane (Lelek et al. 2021).

N-glycans are highly flexible and structurally heterogeneous, making them largely invisible in both high-resolution imaging and traditional structural biology. To bridge this gap and provide a structural basis for our imaging findings, we used GlycoSHIELD (Tsai et al. 2024), a computational tool that uses 1 µs-long Molecular Dynamics (MD) simulations to predict the shielding effect and physical volume occupied by the glycan canopy. By mapping the 3D impact area of the β1-subunit glycans, we aimed to determine how their steric occupancy influences the inter-pump distances and clustering patterns observed in our SMLM data.

In this study, we hypothesized that glycosylation of the Na,K-ATPase β1-subunit not only affects the enzyme’s trafficking but also influences its clustering and organization within the plasma membrane.

## Results

### *In-silico* predictions indicate a shielding effect of the glycans on Na,K-ATPase-β1 subunit

To investigate the extent to which Na,K-ATPase β1 glycosylation masks the protein surface, we used GlycoSHIELD to generate a conformational ensemble of the full Na,K-ATPase complex. We tested the shielding effect against a 10 Å probe, designed to simulate typical protein-protein interactions, across a range of glycan complexities. Three representative glycan conformations (ranging from simple to complex) were selected to recapitulate the impact of glycan heterogeneity (Figure 1a, i-iii). Mapping the quantified shielding onto the protein surface revealed that glycans primarily protect the surface core of the complex (indicated in magenta, shielding score ≥40), while the periphery of the protein surface remains largely unshielded and solvent accessible (represented in gray-cyan Figure 1b, i-iii). Furthermore, sequence-based quantification identified specific residues where shielding peaks occurred, color coded by the specific glycan (*Figure 1c, i-iii*). These data demonstrate that key shielding sites are concentrated on the core surface, notably, both the magnitude and frequency of these peaks increase with glycan complexity, suggesting that more branched structures provide a stronger shielding across a larger surface area.

**Figure 1:**
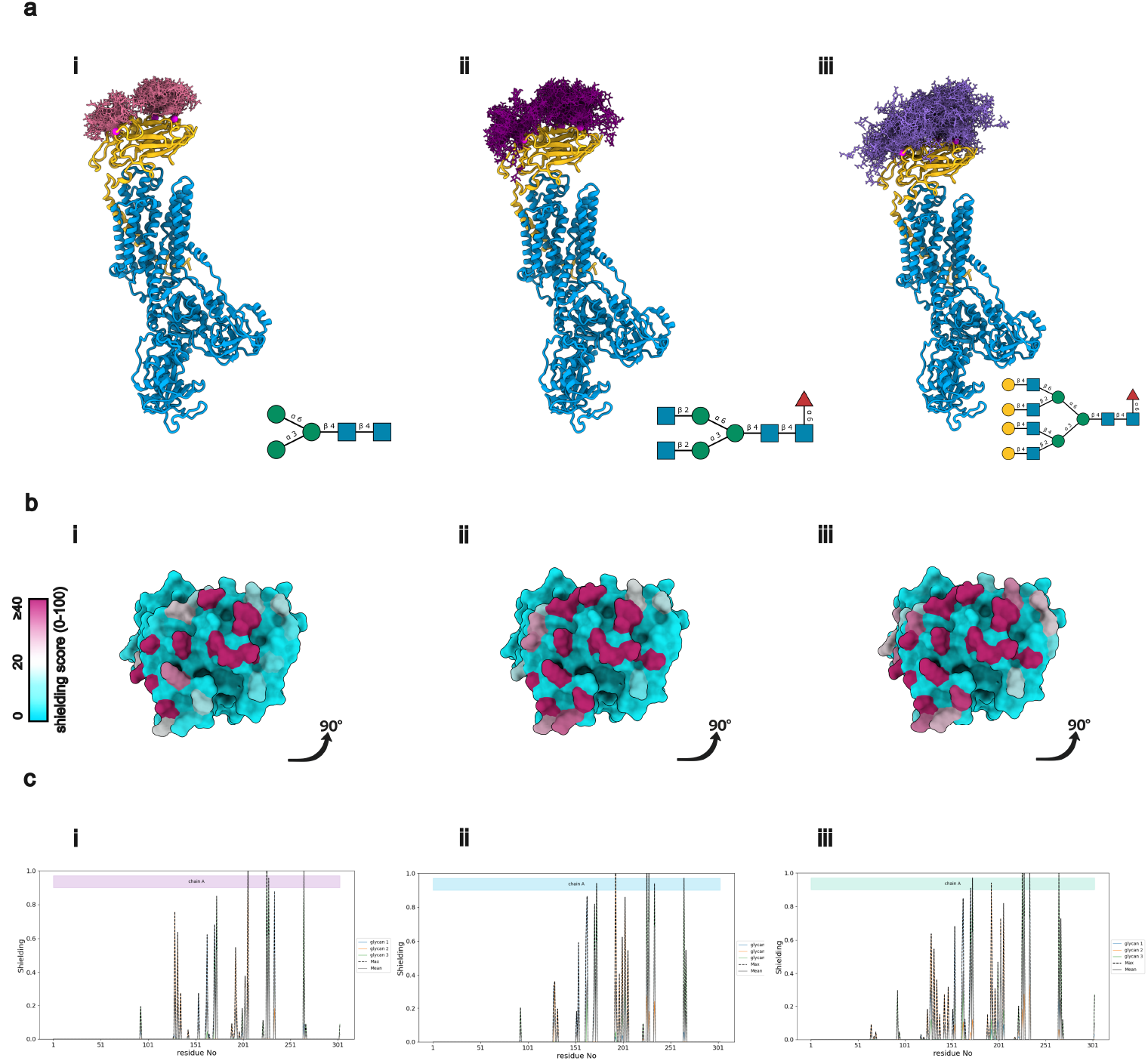
GlycoSHIELD analysis shows that glycans have a shielding effect on the surface of the Na,K-ATPase β1 subunit. a) Structural representation of Na,K-ATPase α1 (blue) and Na,K-ATPase β1 (yellow) with different glycan conformations on three sites, representing three different glycan conformations from less to more complex, (i) Man(a1-3)[Man(a1-6)]Man(b1-4)GlcNAc(b1-4)GlcNAc, (ii) GlcNAc(b1-2)Man(a1-3)[GlcNAc(b1-2)Man(a1-6)]Man(b1-4)GlcNAc(b1-4)[Fuc(a1-6)]GlcNAc, and (iii) Gal(b1-4)GlcNAc(b1-2)[Gal(b1-4)GlcNAc(b1-4)]Man(a1-3)[Gal(b1-4)GlcNAc(b1-2)[Gal(b1-4)GlcNAc(b1-6)]Man(a1-6)]Man(b1-4)GlcNAc(b1-4)[Fuc(a1-6)]GlcNAc. Three different glycan conformations were analyzed to account for large heterogeneity of glycosylation confirmations that can be found on Na,K-ATPase-β1 representing three different glycan lengths and complexities b) Top view of the structural representation of Na,K-ATPase β1 color coded by the shielding effect of the associated glycans indicate that in all three cases the shielding effect is high on the core surface of the protein, while the periphery of the protein surface remains largely unshielded by the glycans as indicated by the shielding score. c) Shielding score along the protein sequence using the 10 Å probe shows distinct shielding peaks along the core surface of the protein indicating that a more complex glycan has a higher shielding effect on a larger surface of the protein, supported by higher and more peaks in (ii) compared to (iii).

### Decreased Surface Localization and Cluster Size of Sodium Pump in Glycosylation-Deficient Mutants Revealed by DNA-PAINT SMLM

To experimentally validate the role of β1 glycosylation in Na,K-ATPase spatial organization, we generated a glycosylation-deficient mutant (3NQ) by substituting the three asparagine residues at positions 158, 193, and 265 with glutamine (*Figure 2a*). Successful mutagenesis and subsequent protein expression were confirmed via DNA sequencing and SDS-PAGE. The 3NQ-GFP mutant exhibited a distinct downward band shift compared to the wild-type (WT) β1-GFP, confirming a reduction in molecular weight consistent with the loss of N-glycans (*Figure 2b*).

**Figure 2:**
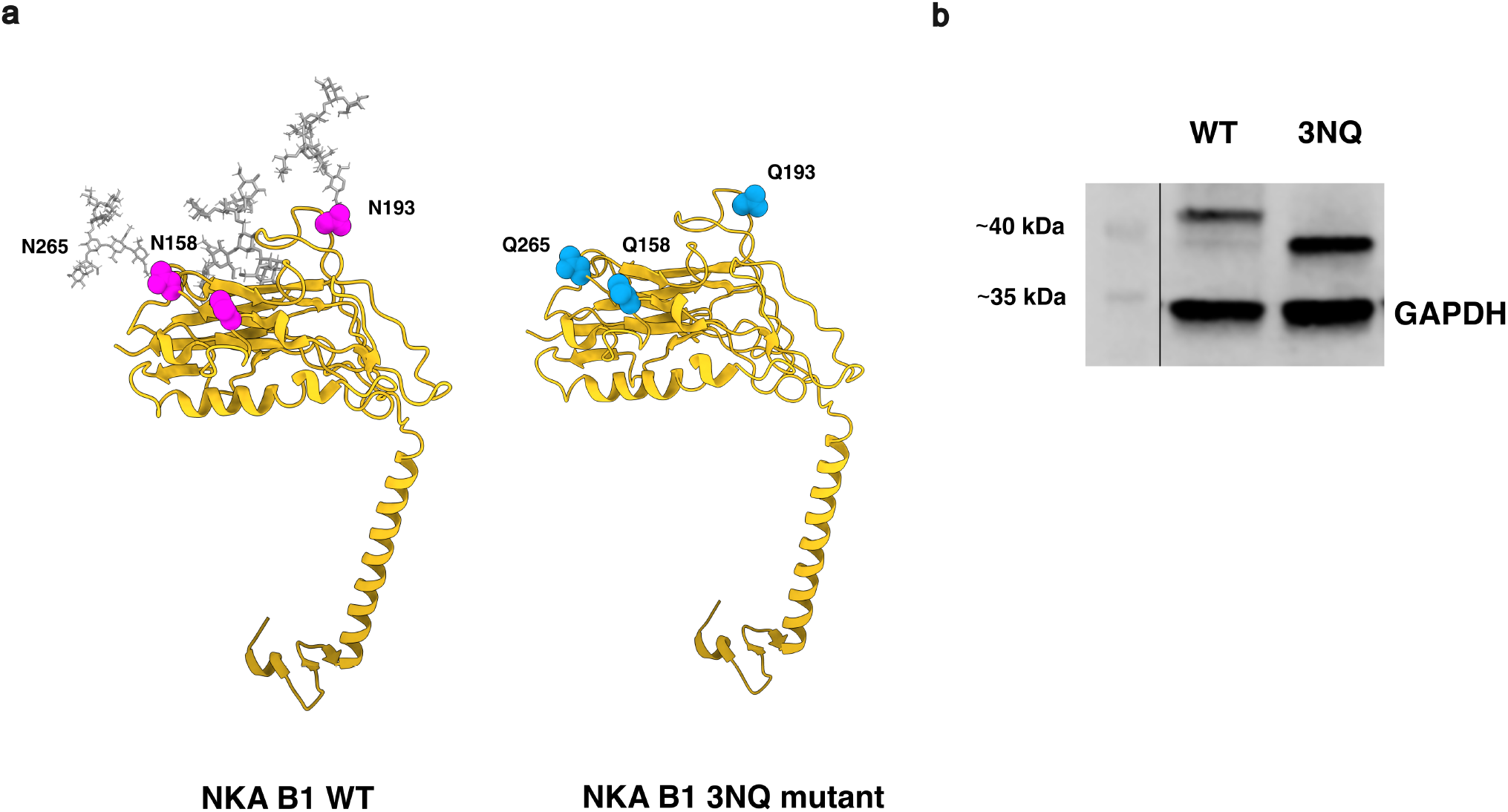
Na,K-ATPase β1 was mutated to inhibit glycosylation at three N sites and generate a 3NQ mutant. a) Structural representation shows wild type Na,K-ATPase β1 (left, PDB 7E1Z) with three N-linked glycosylation sites (magenta, N158, N193, and N256). Na,K-ATPase β1-GFP plasmid was altered with three-point mutations changing asparagine (N) to glutamine (Q) to inhibit glycosylation at three sites (cyan, Q158, Q193, and Q256). b) Western blot of cell lysates transfected with either WT Na,K-ATPase β1 GFP or the mutant 3NQ Na,K-ATPase β1 GFP. Using an anti-GFP antibody, the two conditions show a distinct shift in bands, where the 3NQ most prominent band is lighter, suggesting that the protein complex is not glycosylated, confirming the target mutations.

To resolve the nanoscale organization of the Na,K-ATPase on the plasma membrane, A498 cells expressing either WT or 3NQ constructs were imaged using DNA-PAINT SMLM. Images were acquired using TIRF illumination, effectively isolating the signal to within 100 to 200 nm above the coverslip to reduce background noise and focus the analysis to the proteins expressed on the plasma membrane (*Figure 3a*). Spatial distribution was first assessed using Ripley’s H-function on 3 × 3 µm regions of interest (ROIs). Both conditions showed a non-random distribution, with H-function peaks occurring around 60 nm, indicating a baseline tendency for clustering (*Figure 3b*).

**Figure 3:**
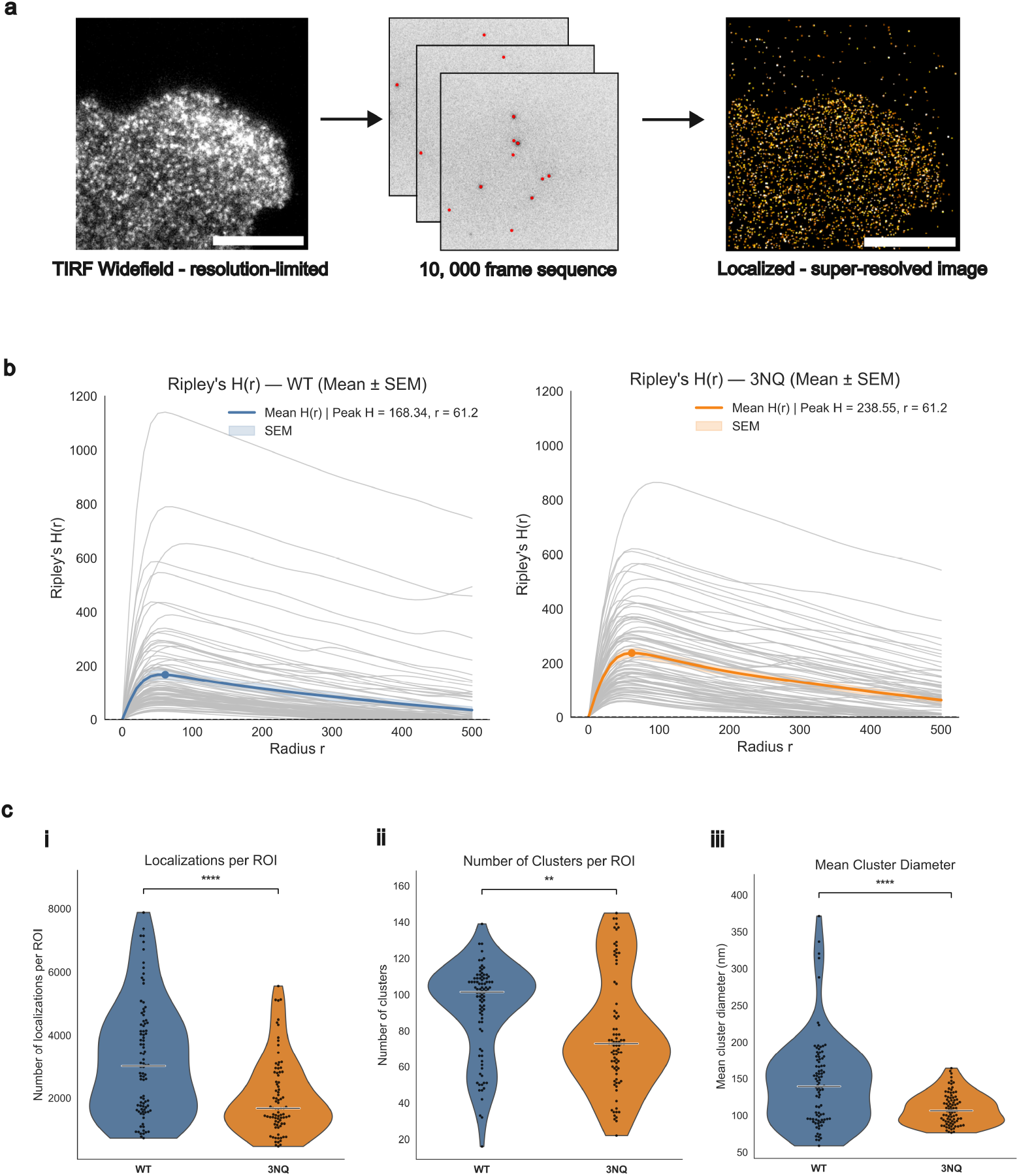
DNA-PAINT SMLM shows that Na,K-ATPase β1 3NQ mutants cluster less frequently and form smaller clusters compared to the WT protein. a) Schematic representation of the SMLM workflow, starting with focusing on the suitable region of the cell in TIRF mode using the GFP signal of the expressed protein (either WT β1-GFP or 3NQ β1-GFP), followed by recording 10, 000 frames of the blinking sequence of ATTO 655 imagers hybridizing to a docking strand on the anti-GFP nanobody, which allows for indirect detection of Na,K-ATPase β1 WT or 3NQ mutant, scale bars 10 µm. b) Ripley’s H curves for selected ROIs for Na,K-ATPase B1 WT (blue) and 3NQ (orange), show that the peak clustering occurs in the mean radius of 61.2 nm. This value was used as epsilon for the DBSCAN analysis. c) (i) 3NQ mutants show approximately two-fold lower number of localizations per ROI as compared to the WT protein, median 1680 vs 3031 points per ROI. DBSCAN analysis (min points= 5, ε = 61.2) shows that 3NQ mutants form a fewer number of clusters median 73 vs 101 for WT (ii) and clusters of a smaller diameter mean 109 nm vs 144 nm for WT (iii). All p < 0.05.

Subsequent analysis using the DBSCAN clustering algorithm, with an ε of 61.2 nm derived from Ripley’s H function and the minimum points for a cluster set to 5, revealed significant differences in membrane organization between the two conditions. WT Na,K-ATPase had a nearly two-fold higher density of localizations compared to the 3NQ mutant, with WT reaching a median of 3031 localizations per ROI compared to 1680 localizations for the 3NQ mutant (*Figure 3c, i*). Furthermore, the 3NQ mutation led to a significant reduction in the frequency of discrete clusters, with a median of 73.0 clusters per ROI compared to 101.5 for the WT (p < 0.05). Cluster morphology was affected in a similar manner, WT formed significantly larger aggregates, characterized by a mean diameter of 144 nm, while 3NQ clusters were significantly smaller, with a mean diameter of 109 nm (p < 0.05, Figure 3c, ii-iii). Taken together, these findings demonstrate that N-glycosylation is a critical determinant of Na,K-ATPase surface density and that its absence negatively affect the formation of stable, large-scale enzyme clusters at the plasma membrane.

## Discussion

Our super-resolved DNA-PAINT SMLM data provide direct biophysical evidence that *N*-glycosylation of the Na,K-ATPase β1 subunit is an important component for its nanoscale clustering at the plasma membrane. The observation that wild-type (WT) pumps form significantly more frequent clusters (median 101 vs 73 per ROI) and larger spatial aggregates (144 nm vs 109 nm mean diameter) than the non-glycosylated 3NQ mutants strongly supports the Galectin-Lattice hypothesis. In this model, *N*-glycans act as essential multivalent handles or docking sites for carbohydrate-binding proteins like galectins, which cross-link adjacent pumps into dense structural assemblies. Without these glycans, the 3NQ mutants lack the necessary molecular dockers resulting in smaller, less frequent clusters and lower overall localization density (median 1680 vs 3031 localizations per ROI) observed in our study. Interestingly, despite the apparent discrepancy in localization density, the Ripley’s H analysis shows that both protein variants have the same clustering peak at around 60 nm, showing a similar intrinsic clustering scale. This suggests that cluster size differences do not only stem from differences in localization density.

These clustering dynamics effectively reject the competing steric repulsion hypothesis as the dominant organizing principle. If the highly hydrated and mobile N-glycan chains primarily functioned as molecular spacers that exert an entropic repulsive force to prevent tight packing, we would expect the non-glycosylated 3NQ mutants to aggregate more densely and form larger clusters than the WT pumps. Instead, our data demonstrates the opposite. The presence of glycans promotes aggregation, proving that the attractive cross-linking forces of the galectin lattice overcome any localized steric hindrance imposed by the sugar chains.

Our *in silico* GlycoSHIELD analyses give a structural context for how the galectin-based clustering occurs without obstructing the pump’s core functional interfaces. The modeling demonstrates that *N*-glycans give a strong physical shielding effect over the core surface of the *β*1 subunit, but that the periphery of the protein surface remains largely unshielded. This selective shielding allows the complex glycan structures, which provide stronger shielding over larger areas as their complexity increases, to project outward and participate in inter-molecular cross-linking. At the same time, the unshielded peripheral regions remain accessible for interactions with the *α* subunit and the FXYD regulatory subunit.

The glycosylation-dependent nanoscale clustering of the Na,K-ATPase has physiological implications beyond its classical role as an ion transporter. The catalytic pumping activity and the ouabain binding capacity of the Na,K-ATPase remain intact independent of the glycosylation state of the *β* subunit. But the spatial organization of the pump is drastically altered. Proper membrane organization into larger, stable clusters is vital because it impacts the pump’s regulatory interactions, its sequestration into specific signaling-rich microdomains, and its function as a cell adhesion molecule.

Proper glycosylation of the β1 subunit is essential for maintaining strong cell-cell adhesion and stable association with adherents junctions, the galectin lattice likely serves as the structural foundation for epithelial integrity. By enzymatically altering N-glycan branching and thereby modulating the density of available docking sites for the galectin lattice, cells have a dynamic mechanism to tighten or loosen epithelial layers and regulate paracellular permeability without requiring a direct change in the absolute quantity or ion-transport activity of the sodium pumps themselves.

## Materials and Methods

### Site-Directed Mutagenesis

To generate glycosylation-defective mutants, a human ATP1B1-GFP plasmid was modified using the Quick-Change kit (Invitrogen) according to the manufacturer’s instructions. Forward and reverse primers were designed to substitute asparagine residues at positions 158, 193, and 265 with glutamine. The reaction was carried out under the following thermal cycling conditions: an initial incubation at 37°C for 15 minutes, followed by denaturation at 94°C for 2 minutes. The cycling conditions consisted of 15 cycles of denaturation at 94°C for 20 seconds, annealing at 57°C for 30 seconds, and extension at 68°C for 3 minutes. After cycling, a final extension was performed at 68°C for 5 minutes, followed by cooling and storage at 4°C. The reaction mixture was transformed into DH5α cells, and clones were selected based on resistance to kanamycin (50 µg/mL). cDNA of resistant clones was prepared by mini-prep and verified by sequencing to ensure the presence of the required mutations and the absence of off-target mutations within the open reading frame.

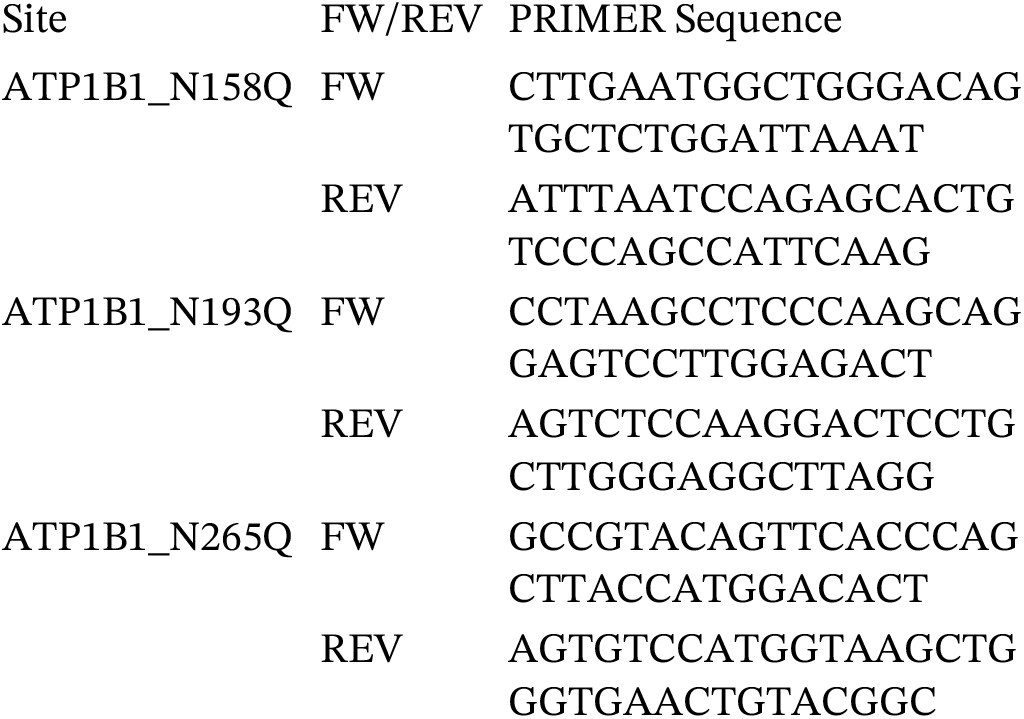

### Western Blot

Transfected cells were washed twice with PBS on ice and treated with a lysis buffer composed of 0.2% Triton-X in PBS supplemented with PhosSTOP (04906845001, Roche) and cOmplete (11697498001, Roche) tablets. Following a 20-minute incubation at -20°C, cells were scraped on ice and the lysate was centrifuged for 20 minutes at 16,000 x g at 4°C. The supernatant was recovered and protein concentration was determination using DC Protein Assay kit (Biorad). Samples were denatured for 10 minutes at 37°C, diluted with 4x Laemmli buffer (161-0747, Biorad) and loaded onto Mini-PROTEAN TGX Stain-Free Gels (4568034, Biorad). Electrophoresis was performed for 45 minutes at 150 V and transferred onto a membrane using a Turbo transfer system (Biorad, 25 A, 25 V, 3 min). The membrane was blocked for 1 hour at room temperature using 5% BSA in TBS-T (0.2% Tween) and incubated overnight at 4°C with rabbit anti-GFP (1:1000, CAB4211, ThermoFischer) and rabbit anti-GAPDH (1:1000, 10494-1-AP, Proteintech) primary antibodies. After three 5-minute washes in TBS-T, the membrane was incubated with anti-Rabbit IRDye 680RD (1:3000, 926-68071, Licor) in 5% BSA TBS Tween for 1 hr at room temperature with mild agitation. Finally, the membrane was washed with TBS three times and imaged using 700 nm excitation with a 2-minute exposure time.

### Cell Transfection and Preparation for Microscopy

The human kidney cancer cell line, A498, was maintained in RPMI medium supplemented with 10% FBS 2 mM L-glutamine, and 1% Penicillin-Streptomycin solution. Cells were seeded in 18-well glass-bottom chambers (iBidi) and transfected after 24 hours using Lipofectamine 3000 for 72 hours as per manufacturer’s instructions. Following transfection, cells were fixed with 4% PFA in PBS for 10 minutes, 3 X PBS wash, permeabilized with 0.5% Triton X-100 for 5 minutes, 3 X PBS wash, and blocked using an antibody incubation buffer (Massive Photonics) for one hour. All steps were performed at room temperature with gentle agitation.

Permeabilized cells were then labelled with MASSIVE-TAG-X2 sdAB anti-GFP (clone: 1H1 & 1B2, 5µM) nanobody at a ratio of 1:200 according to the manufacturer’s instructions, for one hour at room temperature. Samples were washed three times with 1X washing buffer (Massive Photonics), and prior to imaging, immersed in a 1 nM solution of ATTO 655 DS3 imager strands diluted in imaging buffer (Massive Photonics).

### Single-molecule Localization Microscopy (SMLM)

Super-resolution imaging was performed using DNA-Point Accumulation for Imaging in Nanoscale Topography (DNA-PAINT) to resolve the nanoscale spatial distribution of Na,K-ATPase. Single-molecule fluorescence localization data were acquired on a Carl Zeiss Elyra PS.1 microscope equipped with 488 nm and 642 nm excitation lasers using a Plan-Apochromat 100x/1.46 Oil (Zeiss) objective.

Transfected cells expressing the WT β 1-GFP or 3NQ β 1-GFP constructs were identified using the GFP signal, which was excited with a 488 nm laser. To restrict the excitation volume to the plasma membrane and minimize background interference from intracellular fluorophores, the system was operated in Total Internal Reflection Fluorescence (TIRF, q » 66°) mode. The evanescent field extended up to 200 nm into the sample, ensuring that only membrane-bound pumps were selected for subsequent ROI (Region of Interest) identification.

Following the selection of transfected cells, DNA-PAINT imaging was conducted using a 642 nm excitation laser (50% power) and a liquid-cooled EMCCD camera (Andor Technology). To achieve optimal blinking kinetics, an ATTO 655-conjugated DS3 imager strand (∼1 nM) was introduced in specialized imaging buffer. Data were recorded over 10,000 frames with 100 x 100 nm pixel sizes, 80 ms exposure time and an EM gain of 100. Fluorescence emission was isolated using a 650–750 nm bandpass filter (LP650, Zeiss). Data collected represents three biological replicates, where 6-7 images of different cells per condition were recorded.

### SMLM Image Analysis

Following acquisition, raw data were automatically processed for lateral drift correction using ZEN black software. The resulting localization coordinates were filtered to only include high-precision events, excluding localizations with spatial uncertainty > 25 nm or intensities outside the 5th-95th percentile range. To quantify the clustering architecture, 3×3 µm ROIs (5 per recording) were selected, ensuring consistent comparative areas. First Ripley’s H-function was used to determine the degree of non-randomness and identify the characteristic clustering scale, the DBSCAN epsilon (ε) parameter. Then we used the DBSCAN algorithm (minimum points in cluster N=5) to calculate cluster density, mean cluster diameter (maximum distance between localization within a cluster), fraction of points within clusters, and the number of localizations per cluster. Statistical comparisons between WT and 3NQ datasets were performed using the Mann-Whitney U test to account for the non-normal distribution of single molecule data, with significance defined at p < 0.05. Analysis code available at github.com/bruno-stojcic/glyco-PAINT-analysis

### In-silico Glycosylation analysis

The ensemble of glycan trajectories and protein surface shielding of the Na,K-ATPase β1 by glycans were calculated using the GlycoSHIELD web application (dioscuri-biophysics.pages.mpcdf.de/glycoshield-md). The shielding effect was assessed using three distinct glycan structures representing varying complexity levels of ATP1B1 glycosylation, ranging from simple mannose structures to complex branched architectures:

A. Man(a1-3)[Man(a1-6)]Man(b1-4)GlcNAc(b1-4)GlcNAc,
B. GlcNAc(b1-2)Man(a1-3)[GlcNAc(b1-2)Man(a1-6)]Man(b1-4)GlcNAc(b1-4)[Fuc(a1-6)]GlcNAc, and
C. Gal(b1-4)GlcNAc(b1-2)[Gal(b1-4)GlcNAc(b1-4)]Man(a1-3)[Gal(b1-4)GlcNAc(b1-2)[Gal(b1-4)GlcNAc(b1-6)]Man(a1-6)]Man(b1-4)GlcNAc(b1-4)[Fuc(a1-6)]GlcNAc.

A 10 Å probe size was used for the calculations of the protein accessible surface to simulate the steric impact of the glycan shield on protein-protein interactions.

## Acknowledgements

This work was supported by the National Bioinformatics Infrastructure Sweden (NBIS) and ALM, IMT-Stockholm at SciLifeLab and by the National Microscopy Infrastructure, NMI (VR-RFI 2023-00163).

